# Enteric viral infections promote systemic accelerated aging in Drosophila

**DOI:** 10.1101/2025.03.13.643076

**Authors:** Rubén González, Mauro Castelló-Sanjuán, Maria-Carla Saleh

**Affiliations:** Institut Pasteur, Université Paris Cité, Viruses and RNA Interference Unit (Paris, France)

**Keywords:** viral infection, aging, transcriptional aging clocks, RAPToR, Drosophila, pathogenicity

## Abstract

Do viral infections accelerate aging, and does this acceleration scale with pathogenicity? Using transcriptomic aging clocks, we measured biological age in Drosophila infected with four enteric RNA viruses spanning a broad pathogenicity range (i.e. reduction of host lifespan). All pathogenic infections accelerated aging and the magnitude of acceleration tracked pathogenicity. This pattern held across oral and systemic infection routes and was conserved in *Caenorhabditis elegans* where the non-pathogenic Orsay virus produced negligible aging acceleration. Pathway analysis indicated a systemic impact across aging hallmarks with virus- and tissue-specific signatures. Acceleration was comparable in females and males, but host context modulated the acceleration: the bacterial symbiont *Wolbachia* mitigated the virus-induced aging. Notably, biological age remained elevated even after viral clearance. These results demonstrate viruses act as age-distorters and link infection severity to lasting aging consequences, providing a quantitative framework for predicting long-term health effects of viral disease.

## INTRODUCTION

Aging is a complex biological process characterized by progressive functional decline and increased vulnerability to disease and death (Harman 1981). While chronological age measures time since birth, biological age reflects the functional status of an organism and can deviate significantly from chronological age depending on genetic, environmental, and pathological factors (Levine 2013, Khan et al. 2017, Jazwinski and Kim 2019, Zampino et al. 2022). Understanding the mechanisms that accelerate or decelerate aging has profound implications for health and longevity.

Among the various factors that may influence aging, viral infections have emerged as potential modulators of the aging process. Infections with human cytomegalovirus, SARS-CoV-2, HIV, and hepatitis viruses drive premature aging through sustained inflammation, immune system exhaustion, and persistent cellular stress responses (Mekker et al. 2012, Tachtatzis et al. 2015, Lopez Angel 2021, Tsuji et al. 2022). These observations raise fundamental questions about whether viral infections truly accelerate aging processes beyond their direct pathogenic effects.

Quantifying aging acceleration requires robust biological age estimation methods: cellular senescence assays, metabolomic profiling, and DNA methylation patterns (Bafei and Shen 2023, Waaijer et al. 2012, Hertel et al. 2016, Petkovich et al. 2017; Horvath and Raj 2018) allow for estimation of various hallmarks of aging including genomic instability, telomere attrition, cellular senescence, and metabolic dysfunction (López-Otín et al. 2023; Aging Biomarker Consortium et al. 2023). Currently, state-of-the-art computational methods combined with omics data enable the construction of “aging clocks” that capture the molecular complexity of aging across multiple biological layers (Rutledge et al. 2022). These clocks leverage machine learning approaches to integrate thousands of molecular features into composite measures of biological age, with different omics modalities (methylation, transcriptomics, proteomics) often capturing distinct and complementary aspects of aging biology. Transcriptional aging clocks, in particular, leverage genome-wide expression data to capture aging signatures and estimate biological age (Meyer and Schumacher 2021, Bulteau and Francesconi 2022, Jung et al. 2023).

The fruit fly *Drosophila melanogaster* is an ideal model organism for studying aging. Flies have a short lifespan, minimal ethical research concerns, abundant research resources available, and share a large number of functional homologies with humans (Xu and Cherry 2014, Hales et al. 2015, Ugur et al 2016, Kiani et al 2022, Lu et al 2023, Öztürk-Çolak et al 2024). Specifically, the evolutionary conservation of aging mechanisms between flies and humans extends to over 70% of genes associated with aging-related diseases (Piper and Partridge 2018, Tsurumi and Li 2020). In addition, fly aging involves multiple hallmarks conserved with humans such as genomic instability, telomere dysfunction, epigenetic alterations, loss of proteostasis, dysbiosis, mitochondrial dysfunction, and stem cell exhaustion (Clark et al. 2015, Santos and Cochemé 2024).

At the tissue level, fly organs also share similarities with mammals with the fly intestine emerging as a particularly critical determinant of fly lifespan. Indeed, Drosophila intestinal infection and pathology serve as an established model for human intestinal diseases and inflammatory conditions (Apidianakis and Rahme, 2011). Aged intestines progressively lose barrier function, develop dysbiosis, and exhibit stem cell overproliferation. These changes precede death and correlate strongly with survival (dos Santos and Cochemé 2024, Jasper 2020, Biteau et al. 2008; Rera et al. 2012; Regan et al. 2016, Salazar et al. 2023).

Viral infections in Drosophila can elicit phenotypes characteristic of uninfected aged flies, including intestinal stem-cell hyperproliferation and barrier dysfunction (Nigg et al., 2024; Franchet et al., 2025). Because wild flies naturally harbor diverse RNA viruses with varying virulence (Webster et al., 2015; Wallace and Obbard, 2024), we focused on four enteric positive-sense single-stranded RNA viruses—Bloomfield virus, Drosophila A virus (DAV), Drosophila C virus (DCV), and Nora virus—to test whether infection accelerates aging and whether the magnitude of this effect scales with pathogenicity. These viruses establish persistent fecal-oral infections from larval stages into adulthood, initially targeting the intestine before spreading systemically. We used transcriptional aging clocks to quantify biological age and to identify mechanisms and host factors that drive infection-induced age acceleration.

## RESULTS

### Persistent viral infections reduce fly lifespan and induce distinct transcriptional responses

As a system for studying infection-aging relationships, we used persistently infected populations of *Drosophila melanogaster*. These infections were established through oral acquisition of the virus during the larval stage from virus-contaminated environments, simulating natural environmental exposure patterns. Infection occurs upon larval hatching, with virtually all individuals maintaining persistent infections throughout their development and adult lifespan (Castelló-Sanjuán et al. 2025). This approach captures the chronic, long-term effects of persistent infection rather than acute pathological responses.

Lifespan analysis of infected *Drosophila* revealed virus-specific mortality patterns (Fig. 1A). Uninfected flies had a median lifespan of 55.5 ± 12.7 days post adult eclosion (dpe; median ± SD), while infected flies showed significantly reduced lifespans. The most severe effects occurred with DAV (25.0 ± 7.2 dpe) and DCV (24.0 ± 17.1 dpe), representing approximately 55% reduction compared to controls. Bloomfield virus caused intermediate mortality (33.0 ± 11.9 dpe, 40% reduction), while Nora virus showed the mildest impact (46.5 ± 17.9 dpe, 16% reduction).

**Fig 1.**
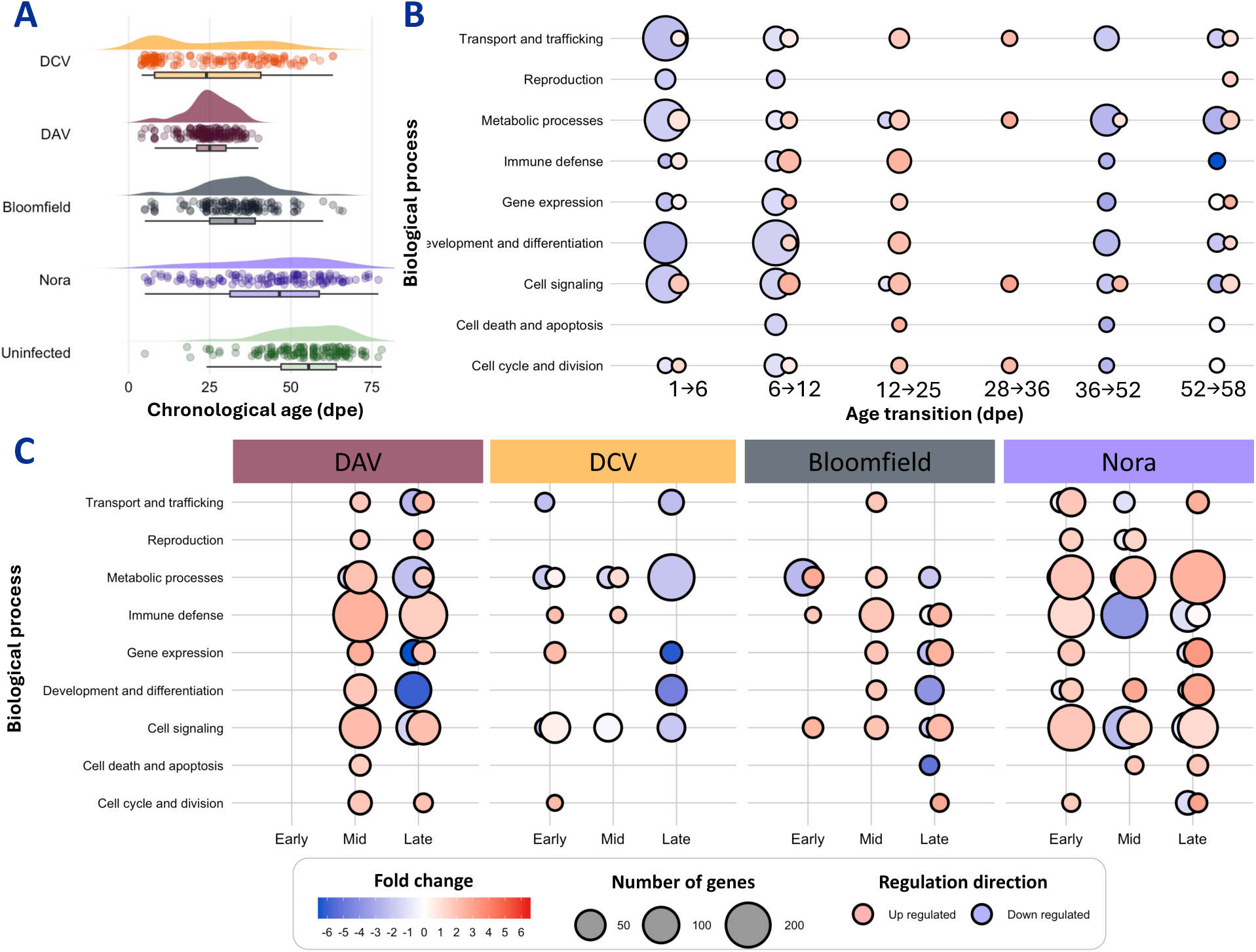
Characterization of viral pathogenicity and associated transcriptional responses. **(A)** Persistent viral infections reduce fly lifespan. Half-violin plot shows the distribution of flies lifespan, with each dot representing the day of death of individual flies and box plots summarizing the data per condition. **(B)** Aging in uninfected flies involves coordinated transcriptional changes across multiple biological processes. Ciruclar symbols show the regulation of biological processes during consecutive age transitions in uninfected flies, comparing early to late infection timepoints (6 vs 1, 12 vs 6, 25 vs 12, and 52 vs 36 dpe). Red indicates upregulation, and blue indicates downregulation. Symbol size represents the number of genes affected, and color intensity indicates the magnitude of fold change. **(C)** Viral infections induce distinct transcriptional responses with virus-specific patterns. Circular symbols show transcriptional regulation in virus-infected flies compared to age-matched uninfected controls at three infection stages. Each virus shows unique temporal patterns of transcriptional regulation across infection stages. Early: 1 dpe, Mid: 12 dpe, Late corresponds to 50% mortality timing for each virus: 25 dpe for DAV, 28 dpe for DCV, 36 dpe for Bloomfield virus, and 52 dpe for Nora virus. Symbol conventions as in panel B.

To understand the molecular basis of these differences in lifespan, we characterized transcriptional changes across time in uninfected and infected flies. Analysis of uninfected flies across six age transitions revealed coordinated expression changes in multiple biological processes (Fig. 1B, Supplementary File 1). Early infection (1-6 dpe) involved moderate transcriptional changes affecting metabolic processes and development. Mid-life infection (6-12 and 12-25 dpe) showed extensive transcriptional remodeling, with upregulation of immune defense pathways and regulation of metabolic and signaling processes. Late infection (36-52 and 52-58 dpe) was characterized by declining expression of genes involved in metabolism, development, and cellular maintenance.

Viral infections induced distinct transcriptional responses that were virus-specific (Fig. 1C, Supplementary File 1). DCV and Nora virus caused widespread transcriptional changes from early to late infection, while DAV and Bloomfield virus elicited more moderate responses. Both highly lethal DAV and mildly lethal Nora virus infections induced extensive transcriptional changes, but they affected different biological processes and followed different temporal patterns. This observation suggested that the relationship between transcriptional responses and aging acceleration might be more complex than simple magnitude of gene expression changes, motivating our subsequent quantitative analysis and the use of computational aging clocks to directly measure biological age acceleration rather than relying on transcriptional similarity as a proxy for aging.

### Transcriptional aging clocks reveal virus-induced aging acceleration

To investigate whether the reduced lifespan was accompanied by acceleration of aging, we employed RAPToR^9^ (Bulteau and Francesconi 2022) to calculate the biological age from gene expression data (Castelló-Sanjuán et al. 2025).

The existing Drosophila aging reference accompanying RAPToR had significant limitations for our study. Most critically, the available reference relies on a two-decade-old microarray dataset (Pletcher et al. 2002) prepared from flies of unknown viral infection status, creating data compatibility issues with modern RNA-seq outputs, and potentially introducing viral contamination artifacts. We therefore constructed a custom aging reference optimized for our experimental conditions. This approach provided critical advantages: confirmed viral infection-free reference samples, RNA-seq data compatibility (avoiding microarray-to-RNA-seq conversion artifacts), higher temporal resolution, and matched experimental conditions. These factors collectively optimized RAPToR performance. We used whole-genome transcriptional data from RNA-seq confirmed virus-free flies collected at eight time points (1, 6, 12, 25, 28, 36, 52, and 58 dpe) from our previous work (Castelló-Sanjuán et al, 2025). Importantly, RAPToR’s interpolation capabilities allow the method to reliably estimate biological ages both within and beyond the original reference time points (Bulteau and Francesconi 2022).

The resulting aging reference accurately captured uninfected fly aging dynamics. When applied to the same uninfected flies used for reference construction, biological age estimates showed strong correlation with chronological age (Pearson’s r = 0.997, *P* < 0.001) (Supplementary Fig. 1A). Leave-one-out cross-validation, when predicting the age of independent samples not used in reference construction, confirmed the accuracy and generalizability of our aging clock (Pearson’s r = 0.935, *P* < 0.001) (Supplementary Fig. 1B).

With this validated aging clock, we assessed the biological age of virus-infected animals at three time points: early stage at 1 dpe, intermediate stage at 12 dpe, and a later stage corresponding to 50% mortality of the host population (25 dpe for DAV, 28 dpe for DCV, 36 dpe for Bloomfield virus, and 52 dpe for Nora virus) (Fig. 2, Supplementary Fig. 2). At 1 dpe, the biological age of uninfected animals was 1.2 ± 0.4 dpe (mean ± SD). Flies infected with Nora virus did not show a significant increase in aging (1.6 ± 0.2 d), but other viruses exhibited accelerated aging even at this early stage: 2.36 ± 0.2 dpe for Bloomfield virus, 1.8 ± 0.3 dpe for DAV, and 2.0 ± 0.2 dpe for DCV. At 12 dpe, the mean biological age of uninfected flies remained similar to their chronological age (12.8 ± 0.9 d), whereas all viral infections significantly increased the biological age: 31.5 ± 3.7 dpe for DAV, 17.8 ± 8.1 dpe for DCV, 24.6 ± 0.3 dpe for Bloomfield, and 25.2 ± 0.6 dpe for Nora. At later time points, DAV-infected flies had a biological age of 54.3 ± 0.1 dpe at 25 dpe and DCV-infected flies had a biological age of 49.6 ± 10.4 dpe at 28 dpe. For the other viruses, there was no accelerated aging at later time points; Bloomfield virus-infected flies showed a reduced but non-significant difference in biological age (31.9 ± 3.9 dpe at 36 dpe), and Nora virus-infected flies exhibited a significant slowing of aging (biological age of 37.1 ± 11.5 dpe at 53 dpe). These results demonstrate that enteric viral infections accelerate aging, with the magnitude and trajectory of aging acceleration displaying virus-specificity.

**Fig. 2.**
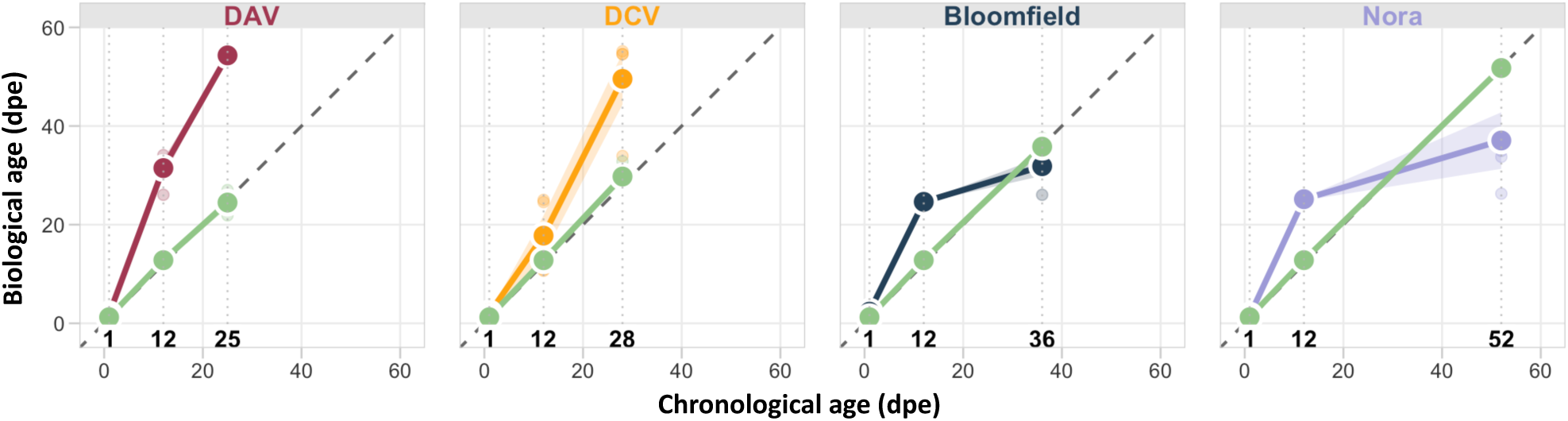
Viral infections disrupts fly aging. Panels show RAPToR-estimated biological age (Y-axis) plotted against chronological age (X-axis) for four different viral infections compared to uninfected controls. Time points include an early stage at 1 dpe, an intermediate stage at 12 dpe, and later stages corresponding to 50% mortality of the host population (25 dpe for DAV, 28 dpe for DCV, 36 dpe for Bloomfield virus, and 52 dpe for Nora virus). Colored lines represent virus-infected flies (DAV in red, DCV in orange, Bloomfield virus in blue, Nora virus in purple), while green lines represent uninfected controls. Large points represent the mean biological age of four biological replicates per condition, and shaded areas represent standard error. The dashed diagonal line indicates perfect correlation between chronological and biological age

### Virus-induced accelerated aging is systemic, temporally dynamic, and targets aging pathways and tissues in a virus-specific manner

We investigated whether the observed aging acceleration reflects broad systemic effects (affecting multiple tissues and pathways) or arises from modulation of specific pathways. To do so, we created specialized aging references using genes associated with key aging processes (such as telomere shortening, stem cell dysfunction, and protein damage) and major tissues (gut, fat body, muscle, brain, and ovary). We then assessed acceleration of individual aging processes by estimating biological age to chronological age ratios for each process using our specialized references at the three critical timepoints described in the previous section: early, mid, and late infection. Aging pathways were established from proved aging hallmarks (López-Otín et al. 2023) and curated from FlyBase pathways, and tissue-specific genes were obtained from FlyBase tissue enrichment data (Öztürk-Çolak et al. 2024). Complete gene lists and identifiers can be found in Supplementary File 2.

Temporal analysis revealed distinct patterns of virus-induced aging acceleration that diverge from normal aging trajectories (Fig. 3). Uninfected flies maintained median biological to chronological age ratios close to 1 across all timepoints and tissue-specific references, demonstrating that our aging references accurately estimated aging in all organs and that hallmarks of aging pathways alone could reliably predict biological age. This established the baseline for normal aging progression. In contrast, viral infections accelerated aging from the very first day of adult life (early infection timepoint), with most virus-pathway combinations showing elevated aging ratios immediately upon adult eclosion. This acceleration was maintained through the mid-infection timepoint (12 dpe), indicating sustained aging pressure during active viral infection phases. Late infection timepoints revealed divergence between viruses. DAV and DCV maintained a high aging acceleration into late infection, with many pathways showing continued accelerated aging effects. Bloomfield virus showed broad pathway targeting that decreased over time, with aging pace returning to the levels seen in uninfected flies, indicating resolution of aging effects despite continued infection. However, Nora virus showed a pattern where aging effects were broad only at the mid-timepoint, with late-stage aging falling below the biological to chronical age ratio of uninfected controls, suggesting that infected flies that survived until this late stage had deployed tolerance mechanisms that slowed their aging below normal rates.

**Fig. 3.**
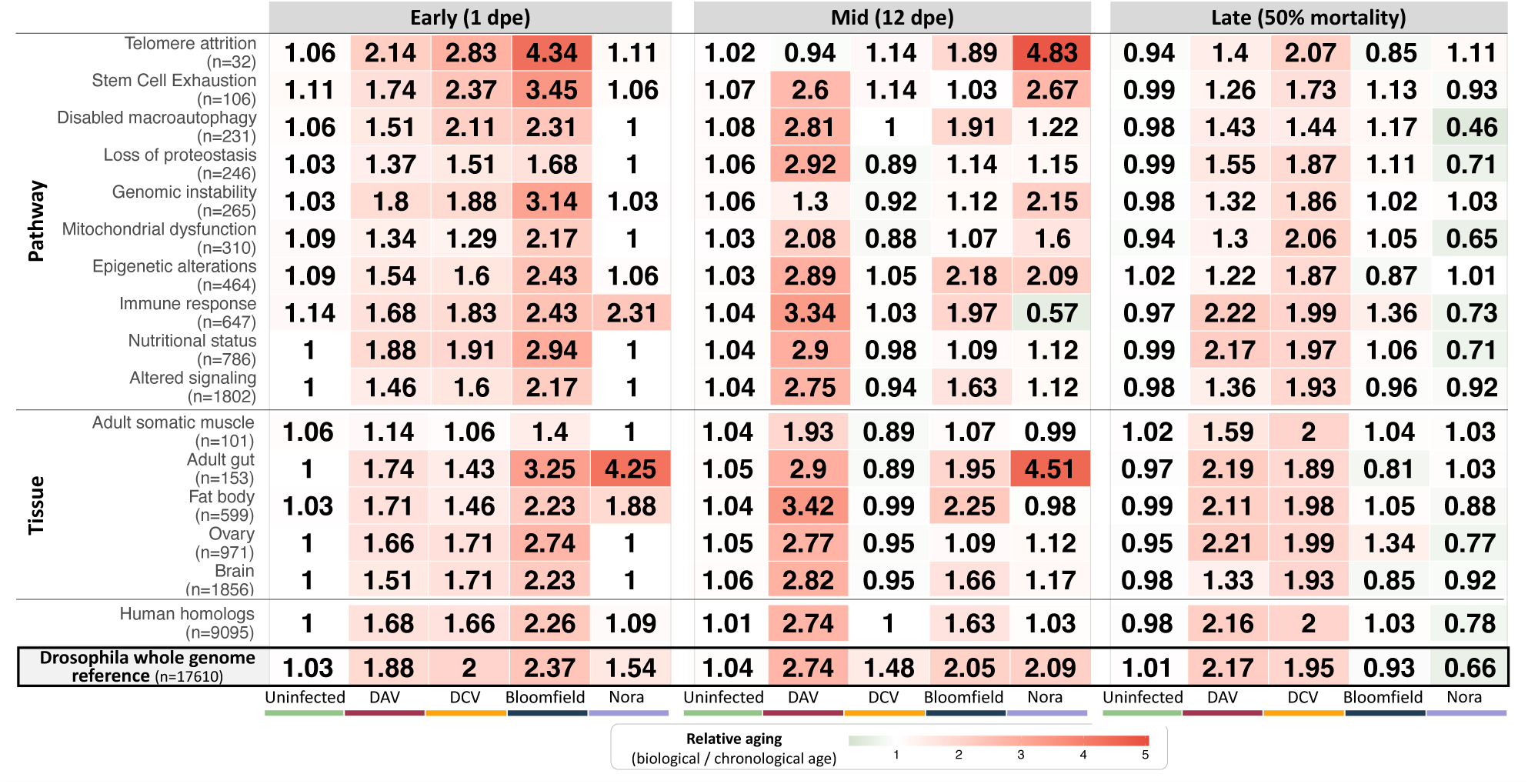
Temporal dynamics of virus-specific targeting of aging pathways and tissues. Heatmap shows median relative aging acceleration (biological age/chronological age) across infection progression for tissue-specific and pathway-specific aging references. Early infection (1 dpe), mid-infection (12 dpe), and late infection timepoints (DAV: 25 dpe, DCV: 28 dpe, Bloomfield: 36 dpe, Nora: 52 dpe, representing ∼50% population mortality). For uninfected flies, the late timepoint represents the median across samples from 25, 28, 36, and 52 dpe. Values >1.0 (red) indicate accelerated aging, values <1.0 (green) indicate slowed aging, and values 1.0 (white) indicate normal aging progression. Numbers in parentheses indicate the number of genes in each reference set. Median absolute deviation values provided in Supplementary File 1.

Analysis of tissue-specific aging acceleration revealed that all tissues tested showed evidence of accelerated aging at some timepoints, confirming the systemic nature of virus-induced aging effects. However, the adult gut and fat body emerged as the most critical targets of virus-induced aging across all viruses and timepoints, and in the case of the Nora virus infected individuals, were the only tissues with accelerated aging. Among aging pathways, immune response and telomere attrition showed elevated aging ratios in most conditions and timepoints. Beyond these common targets, individual viruses showed distinct pathway specificity: DAV and DCV exhibited broad pathway targeting across multiple timepoints, Bloomfield virus affected broad pathways that decreased over time, while Nora virus only affected broad pathways at the mid-timepoint. DCV-infected animals at the mid-infection timepoint showed an aging acceleration that exceeded the effects attributable to any individual pathway or tissue studied (whole genome aging ratio of 1.48 versus 0.86 to 1.14 in tissues and pathways respectively). This suggests that at this infection stage, DCV induces aging through different pathways and/or complex interactions among the aging hallmarks we examined.

To assess evolutionary conservation, we created aging references using only Drosophila genes with direct human orthologs. These human-ortholog-only references showed aging acceleration patterns largely concordant with the Drosophila whole genome reference (last two rows Fig. 3), suggesting evolutionary conservation of virus-induced aging mechanisms and indicating the potential of these findings for understanding viral impacts on human aging and longevity.

To confirm that aging acceleration represents truly systemic effects rather than pathway-specific dysregulation, we performed exclusion analysis using aging references constructed from all genes except those associated with each pathway or tissue. When pathway-specific or tissue-specific genes were excluded from the aging references, we still detected comparable virus-induced aging acceleration across all timepoints and conditions. The exception was DCV at the mid-timepoint, where epigenetic alterations and immune response genes were required to detect aging acceleration (Supplementary Fig. 3). This demonstrates that viral aging effects generally extend beyond the dysregulation of any single biological process or tissue, confirming that virus-induced aging represents a broad, systemic acceleration of biological age that cannot be attributed solely to targeted disruption of particular aging pathways.

### Viral load relationships suggest distinct aging mechanisms

To explore if viral loads relate to aging acceleration, we evaluated correlations between virus RNA accumulation (measured by RT-qPCR targeting the viral polymerase) and aging across the infection timeline (Fig. 4). The results showed a different relationship for each virus, suggesting distinct mechanisms of virus-induced aging.

**Fig. 4.**
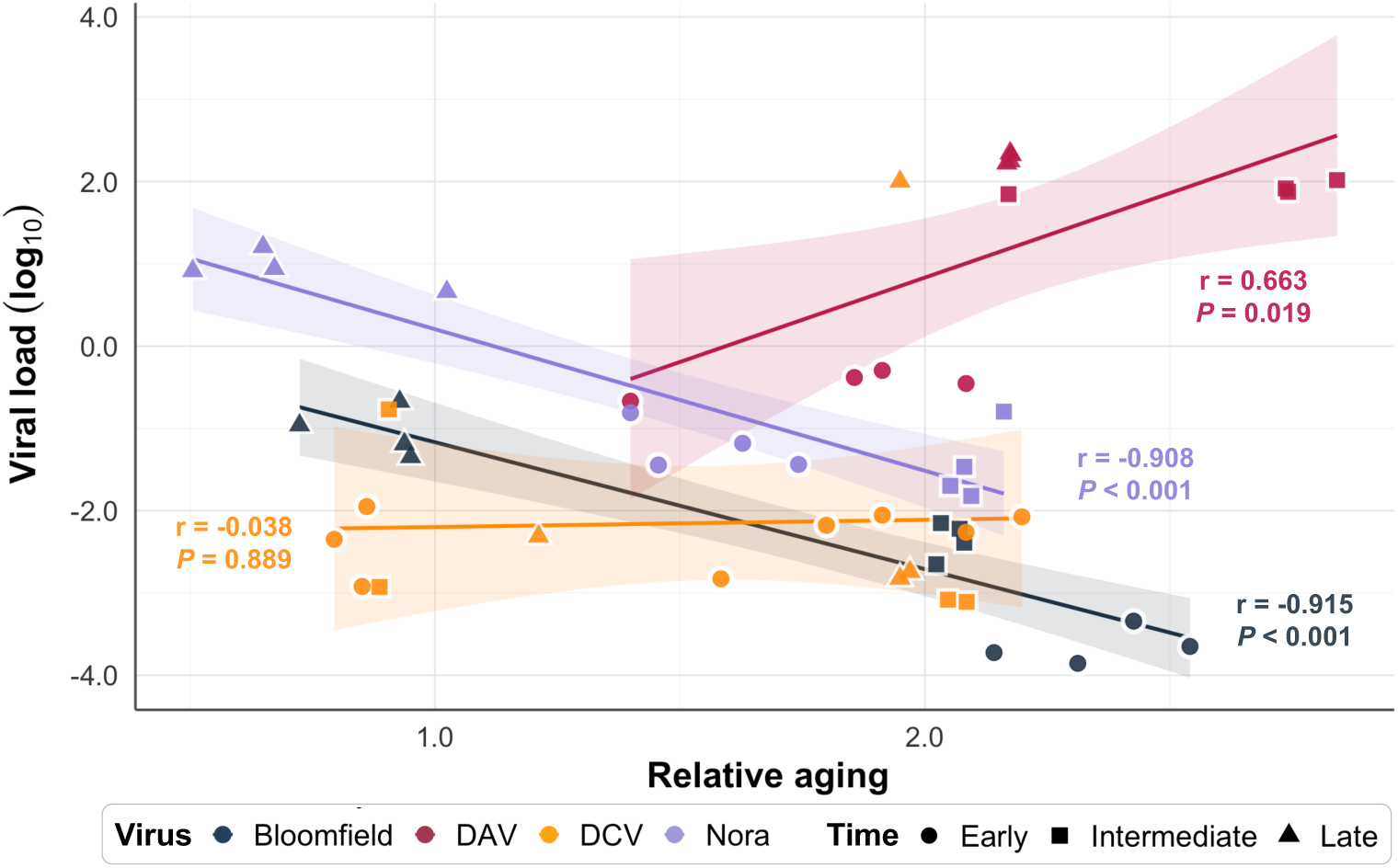
Correlation between relative aging (ratio between the biological and chronological age at a given timepoint) and viral load (viral RNA accumulation as determined by qRT-PCR) for four persistent viral infections in Drosophila. Early time points are 1 dpe (and 6 dpe for DCV), the intermediate time point is 12 dpe, and the late time point corresponds to 50% mortality of the host population. Solid lines represent linear regression fits for each virus. Shaded areas indicate 95% confidence intervals. Pearson correlation coefficients (r) and *P*-values are displayed in color-matched text for each virus.

DCV showed no relationship between virus levels and aging (Pearson’s r = −0.038, *P* = 0.889), indicating that aging acceleration occurs independently of how much virus accumulates in the host. DAV showed a strong positive correlation between virus levels and aging acceleration (r = 0.663, *P* = 0.019). This indicates that viral accumulation directly damages the host, with more virus accumulation causing proportionally more aging. Bloomfield and Nora viruses showed strong inverse relationships: aging acceleration was greatest when virus levels were lowest (Bloomfield virus: r = −0.908, *P* < 0.001; Nora virus: r = −0.908, *P* < 0.001). This pattern suggests that the host’s efforts to fight these viruses drives aging. During periods when the immune response successfully suppresses viral acumulation, the costs of this defense may accelerate aging more than the infection itself.

These findings, combined with the virus-specific pathway targeting described above, demonstrate that virus-induced aging can occur through multiple mechanisms.

### Viral pathogenicity positively correlates with aging acceleration

Having established that less-lethal viruses produce smaller aging effects, we tested our hypothesis that viral pathogenicity (here defined as the reduction in host lifespan relative to uninfected controls) quantitatively predicts aging acceleration.

To establish the low end of the pathogenicity spectrum, we analyzed Orsay virus (OrV) infection in *C. elegans* larvae. OrV is an enteric RNA virus that infects nematodes with minimal pathogenicity (Félix et al 2011; Ashe et al. 2013). Using the transcriptional data of uninfected larvae (Castiglioni et al. 2024) we established an aging reference using the same methodology described above. Using this reference, we observed that OrV infection had minimal impact on nematode aging. The biological age of infected nematodes tracks closely with chronological age except for a brief period of acceleration coinciding with peak viral replication (Supplementary Fig. 4, and Castiglioni et al. 2024). This confirms that minimally-pathogenic viruses cause overall negligible aging acceleration and provides a low-pathogenicity data point for correlation analysis.

To quantify the relationship between pathogenicity and aging effects, we calculated relative pathogenicity (lifespan reduction) and relative aging acceleration using area under the curve calculations (see Methods). Across oral infections (four viruses in Drosophila and OrV in nematodes), we observed a significant positive correlation between infection pathogenicity and accelerated aging (Pearson’s r = 0.902, *P* = 0.036, Fig. 5A). This correlation suggests that aging acceleration is a key mechanism by which enteric viruses reduce host lifespan.

**Figure 5.**
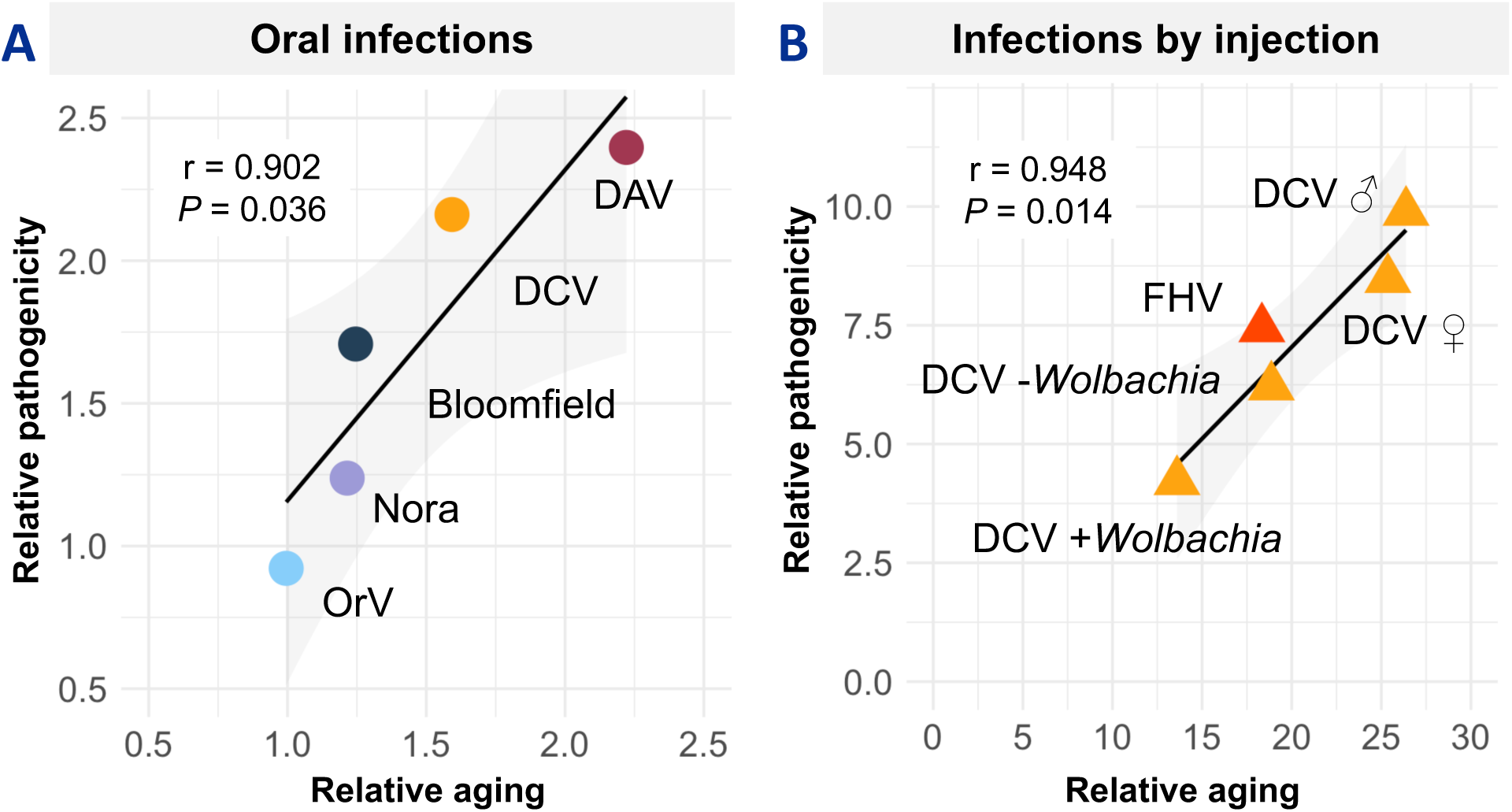
Viral pathogenicity correlates with aging acceleration across infection routes. Correlation between relative aging acceleration (ratio between the biological and chronological age) and relative pathogenicity (ratio between uninfected and infected host lifespan) for **(A)** Oral infections: Four Drosophila viruses spanning a wide pathogenicity range (DAV, DCV, Bloomfield virus, Nora virus) plus non-pathogenic Orsay virus (OrV) in *C. elegans* larvae. Each point represents a different virus. **(B)** Injection infections: Correlation analysis extended to injection-delivered viruses, including DCV in male (♂) and female (♀) flies, Flock House Virus (FHV), and DCV in flies with (+*Wolbachia*) or without (-*Wolbachia*) symbiotic bacteria. Values >1.0 indicate increased mortality or aging acceleration relative to uninfected controls.

To test whether this relationship extends beyond natural oral infections, we examined viral infections initiated by injection. Here, injection bypasses gut barriers and typically results in increased increased viral pathogenicity. Using data from injected DCV (Salminen et al. 2024, Martinez et al 2019) and Flock House Virus (FHV) (Sheffield et al. 2021), we found that aging was higher in injected animals and positively correlated with pathogenicity (Pearson’s r = 0.948, *P* = 0.014, Fig. 5B).

### Host factors modulate virus-induced aging

Having established virus-specific aging mechanisms, we next examined whether host characteristics could influence the magnitude of infection-induced aging. We tested various factors that could modulate virus-induced aging acceleration.

First, we examined host sex differences, as our initial analysis used female flies and *Drosophila* exhibits sex-specific aging pathology and transcriptional and immune responses (Hill-Burns and Clark 2009, Regan et al. 2016, Belmonte et al 2020, Vincent and Dionne 2021). Using a transcriptional dataset that included DCV-infected males and females (Salminen et al. 2024), we found comparable biological ages in uninfected animals of both sexes and similar significant acceleration of aging upon DCV infection (Fig 6A).

**Fig. 6.**
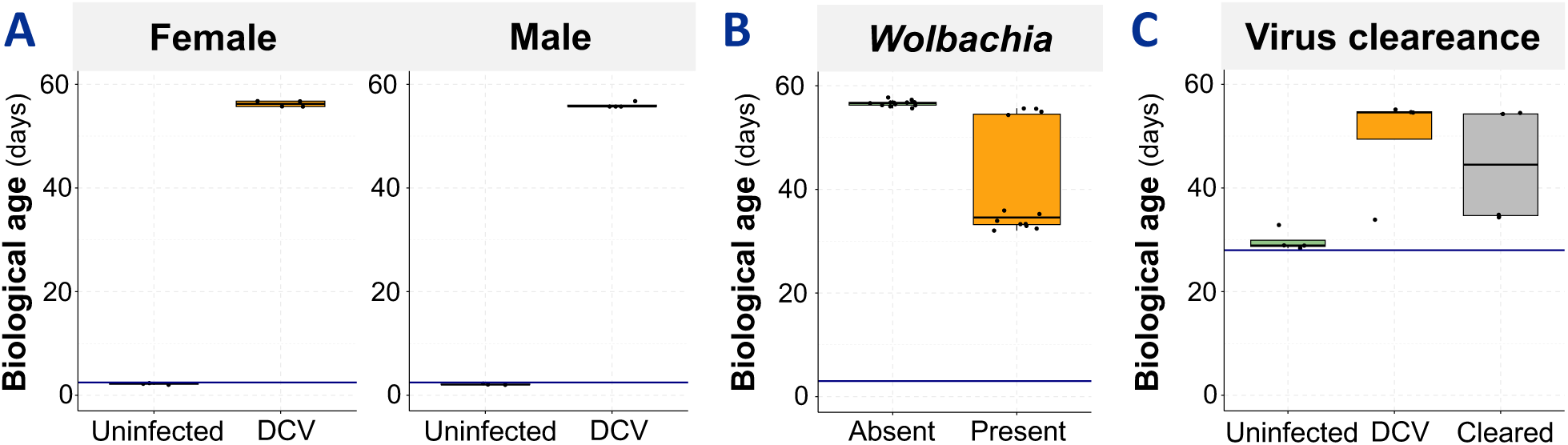
Host factors modulate virus-induced aging. Biological age estimates showing **(A)** sex-independent aging acceleration in DCV-infected flies, **(B)** *Wolbachia*-mediated protection against virus-induced aging, and **(C)** persistent aging effects in flies that cleared viral infection. Blue horizontal lines represent chronological age; boxplots show biological age distributions.

Second, we studied the effect of *Wolbachia*, a common bacterial symbiont of insects known to reduce viral pathogenicity and transmission (Teixeira et al. 2008, Moreira et al. 2009). Using a transcriptional dataset of flies injected with DCV and either harboring or not *Wolbachia* (Martinez et al. 2019), we observed that harboring *Wolbachia* mitigated the aging induced by DCV. At 3 dpe, *Wolbachia*-free flies injected with DCV had a biological age of 56.7 ± 0.6 dpe (median ± SD). In contrast, flies harboring *Wolbachia* had a biological age of 34.6 ± 10.6 dpe (Fig. 6B).

Third, we examined viral clearance. Although viral infections are typically persistent in fly populations, some individuals can clear DCV infection (Mondotte et al. 2018, Castelló-Sanjuán et al. 2025). By examining data from flies persistently infected with DCV (Castelló-Sanjuán et al. 2025), we observed that both the DCV-infected flies and virus-cleared flies showed significantly higher biological age than uninfected flies; at 28 dpe the uninfected flies had a median biological age of 28.9 d, compared to 54.6 ± 10.4 dpe for the DCV-infected flies and 44.5 ± 11.4 dpe for flies that had cleared DCV-infection (Fig. 6C). This suggests that once aging acceleration is triggered by viral infection, it persists even after viral clearance.

## DISCUSSION

Viruses have long been known to modulate their hosts in remarkable ways (Christiaansen et al. 2015, Blanc and Michalakis 2016), but their role in modulating aging process has only recently begun to be explored. Our work demonstrates that viral infections modulate biological age in proportion to their pathogenicity, establishing viruses as quantifiable “age-distorters” that interfere with host aging trajectories (Teulière et al. 2021).

Our findings reveal three distinct strategies by which viruses accelerate aging. DCV acts as an aging “switch” that triggers cellular aging programs at a magnitude that is independent of viral load, potentially explaining why mild chronic infections sometimes have disproportionate long-term consequences. DAV causes dose-dependent aging where higher viral loads produce proportionally greater aging acceleration, indicating direct pathogen-mediated damage. Nora and Bloomfield viruses show inverse correlations where aging peaks during viral suppression by the immune system, demonstrating that host defense responses can drive aging more than the pathogens themselves. Late-stage Nora virus infection slows aging below normal rates, raising the possibility that certain microorganisms could serve as aging mitigators. Pathogenic viral infections have been shown to provide benefits depending on environmental conditions (Xu et al. 2008, González et al. 2020), and pathogenic viruses can evolve into mutualistic relationships in certain environments (González et al. 2021). This opens possibilities for discovering naturally occurring viruses or reshaping existing host-virus interactions to establish beneficial relationships that reduce host aging.

Evolutionary conservation strongly suggests translational relevance. Most aging-associated gene families originated hundreds of millions of years ago (Bonnefous et al. 2024). These ancient pathways represent universal targets, as viral proteins have convergently evolved to directly interact with host aging networks across phylogenetically distant species (Teulière et al. 2023). Consistent with this deep conservation, both universal and species-specific aging mechanisms operate across diverse organisms from flies to mammals (Tyshkovskiy et al. 2023), indicating that viral modulation of aging likely targets both conserved pathways and species-specific mechanisms. We observe consistent acceleration of aging in gut and fat body tissues, which parallels intestinal and metabolic dysfunction in mammalian aging (Funk et al., 2020). In addition, host-associated microorganisms have been shown to modulate viral infections and transmission in diverse hosts (Kane et al 2011, Kuss et al. 2011, Pfeiffer and Virgin 2016). We demonstrate that *Wolbachia*’s protective effects extend to mitigation of virus-induced aging, showing that beneficial microorganisms can modulate infection-induced aging effects.

Methodological robustness enables broad adoption. Our aging clock framework demonstrates consistent performance across different datasets, fly strains, and experimental conditions, aligning with RAPToR’s demonstrated compatibility between datasets even though optimal results are achieved within datasets (Bulteau and Francesconi 2022). Pathway-specific analyses confirm that viruses induce genuine aging perturbation. Hence, our framework and dataset can be widely adopted for aging research in Drosophila and potentially other organisms.

Persistent aging effects require further investigation. Flies that successfully cleared infections retained elevated biological age to varying degrees, indicating that aging acceleration involves fundamental cellular changes beyond transient physiological disruption. Our observations align with findings in humans, where irreversible changes in the epigenome and immune system have been observed following clearance of viral infections (Yates et al. 2021, Hlady et al. 2022). Similarly, clearance of chronic hepatitis C virus only partially reversed aging in some individuals while having no effect in others (Oltmanns et al. 2023). This persistent effect suggests that targeted interventions may be needed beyond simply supporting viral clearance.

Future directions include identifying molecular mechanisms by which different virus target aging pathways and testing whether established anti-aging interventions can prevent or reverse virus-induced aging acceleration. The framework should be extended to DNA viruses, bacteria, and fungi to reveal general principles of microbe-host aging interactions. Clinically, transcriptional aging clocks could complement traditional markers by revealing long-term health impacts not apparent through conventional assessments, particularly for infections causing mild, short term symptoms but substantial aging acceleration. The discovery that some microorganisms can slow aging opens therapeutic possibilities for developing beneficial microorganisms as aging interventions, though such approaches require rigorous safety evaluation.

Our findings on virus-induced aging provide a foundation for investigating aging mechanisms, identifying anti-aging targets, reducing infection pathogenicity, and leveraging viruses as tools to modulate aging.

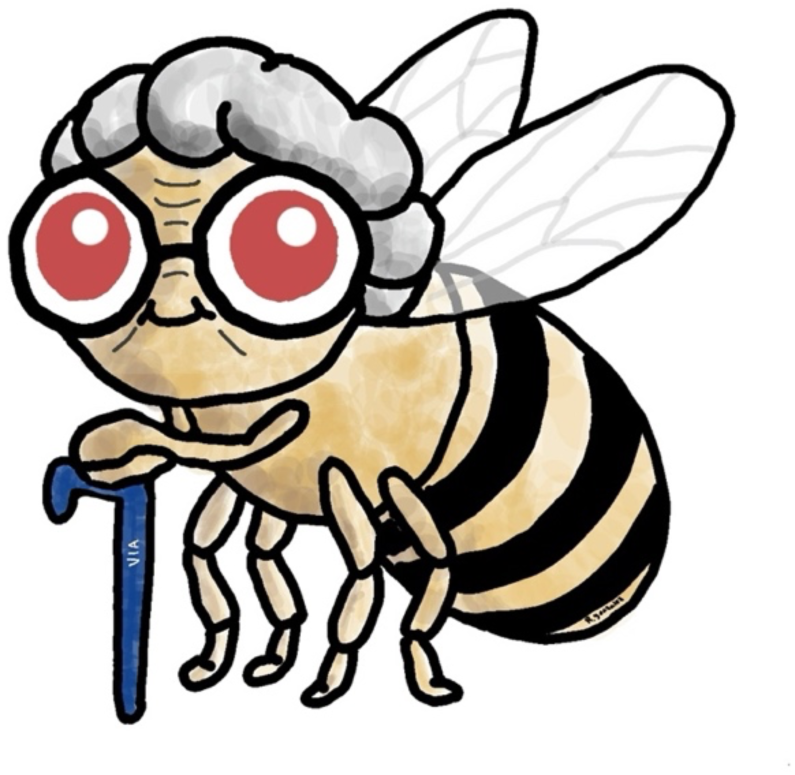

## MATERIAL AND METHODS

### Dataset biological information

All RNA samples come from whole flies or nematodes. Unless otherwise indicated, all flies are *Wolbachia*-free and mated *D. melanogaster*. Persistent infected flies data belong to four pools of 5 *w^1118^* flies per virus and time point (Castelló-Sanjuán et al. 2025). DCV male and female dataset has four pools of 4-7 dpe 10 Oregon-RT virgin flies per condition (Salminen et al. 2024). DCV and *Wolbachia* dataset has 12 pools of 6-9 dpe 10 *w^1118^* flies per condition (Martinez et al. 2019). FHV dataset has four measures per condition of an undetermined number of 6-9 dpe Oregon-R male flies (Sheffield et al. 2021). OrV dataset has 3 pools of 1200 *C. elegans* nematodes per condition (Castiglioni et al. 2024)^11^.

### Transcriptomic data

Persistent infected flies’ raw sequencing reads can be found in the Sequence Read Archive under the BioProject PRJNA1235228. For injected flies raw sequencing reads were retrieved from the same archive: FHV from BioProject PRJNA644593, DCV and *Wolbachia* from BioProject PRJEB21984, DCV in male and female from BioProject PRJNA1084779. The *C. elegans* mapped reads were kindly provided by Santiago Elena’s lab. Raw sequencing reads were mapped to the *D. melanogaster* genome release dmel_r6.56 with STAR (v2.7.11b) (Dobin et al. 2013). Feature counting was done with HTSeq (v0.11.2) (Putri et al. 2022) using the default settings.

### Biological categories

To categorize the biological processes affected during aging and viral infections, we established categories to capture major cellular and physiological processes. Based on GO terms, these categories encompass: Immune defense (GO terms containing keywords such as immune, antimicrobial, antiviral, defense, inflammatory, pathogen, bacteria, virus), Transport & trafficking (transport, localization, secretion, vesicle, trafficking, endocytosis, exocytosis), Cell cycle & division (cell cycle, mitosis, division, proliferation, dna replication, chromosome), Gene expression (transcription, translation, expression, rna, protein, gene regulation, chromatin), Development & differentiation (development, morphogenesis, differentiation, organogenesis, embryonic, larval, growth), Cell signalling (signal, signaling, response, communication, receptor, kinase, phosphorylation), Cell death & apoptosis (death, apoptosis, necrosis, autophagy, programmed cell death), Metabolic processes (metabolic, metabolism, catabolism, anabolism, biosynthetic, glycol, lipid, carbohydrate, amino acid, energy), and Reproduction (reproduction, sex, mating, gamete, oogenesis, spermatogenesis).

### Age estimation

We estimated biological age from transcriptomic data using the RAPToR package (v1.2.0) (Bulteau and Francesconi 2022) in R (v4.3.2) in the RStudio development environment (v2024.04.2+764). First, a reference was created from uninfected flies collected on days 1, 6, 12, 25, 28, 36, 52, and 58 post-eclosion (Castelló-Sanjuán et al. 2025). Gene expression counts were fitted to a generalized additive model via RAPToR’s (v1.2.0) *ge_im* function using a cubic regression spline with eight knots. We then performed a principal component analysis (PCA) with the *prcomp* function in the ‘stats’ package, determining that 29 principal components should be retained (Supplementary Fig. 5) by summing those with a cumulative variance below 0.99 and then adding one. Following RAPToR methodology, we included components with intelligible dynamics. As a control for overfitting, using only the first 10 components provided similar results (Supplementary Fig. 6). This approach leverages RAPToR’s reference interpolation capability, which decomposes the expression matrix into components that summarize gene expression dynamics and interpolates these components with respect to time to reconstruct full interpolated gene expression profiles. The final reference was constructed using RAPToR’s *make_ref* function with 1000 interpolation points. Subsequently, we estimated the biological ages of additional transcriptomic datasets by applying RAPToR’s *ae* function to this reference. In specific cases described in the main text, new references were generated using the same approach but were restricted to subsets of genes of interest (Supplementary Files 1) derived from the original uninfected dataset. For analyses involving virgin fly datasets (Salminen et al. 2024), genes that are differentially expressed after mating (Newell et al. 2020) were excluded from reference construction to avoid mating status confounding effects (Supplementary File 3).

### Pathway and tissue-specific gene sets curation

Aging hallmark genes were curated from established frameworks (López-Otín et al. 2023) and cross-referenced with FlyBase Gene Ontology annotations (release FB2024_02). Pathways were defined using the following GO terms: Genomic Instability -DNA damage response (GO:0006974); Telomere Attrition - Telomere maintenance (GO:0000723); Epigenetic Alterations - Chromatin organization (GO:0006325); Loss of Proteostasis - Response to topologically incorrect protein (GO:0035966), protein ubiquitination (GO:0016567); Disabled Macroautophagy - Autophagy (GO:0006914), protein targeting to vacuole (GO:0006623); Deregulated Nutrient Sensing - Response to insulin (GO:0032868), TOR signaling (GO:0031929), response to starvation (GO:0042594), lipid metabolic process (GO:0006629); Mitochondrial Dysfunction - Mitochondrion organization (GO:0007005), electron transport chain (GO:0022900); Cellular Senescence - Cellular senescence (GO:0090398); Stem Cell Exhaustion - Stem cell population maintenance (GO:0019827); Altered Intercellular Communication - Signaling (GO:0023052); Immune Response - Immune response (GO:0006955), defense response (GO:0006952), defense response to other organism (GO:0098542), response to bacterium (GO:0009617).

Tissue-enriched genes were obtained from FlyBase tissue ontology annotations using the following anatomical terms: Ovary (FBbt:00004865), Adult somatic muscle (FBbt:00058784), Fat body (FBbt:00005066), Adult gut (FBbt:00007513), Brain (FBbt:00005095).

The number of genes per pathway and tissue are indicated in Figure 3, with complete gene lists provided in Supplementary File 2.

### Viral load quantification

Viral loads were measured by RT-qPCR in our previous work (Castelló-Sanjuán et al. 2025). Individual flies were homogenized in TRIzol for RNA extraction, followed by cDNA synthesis using random primers and qPCR using virus-specific primers. Viral RNA levels were normalized to the housekeeping gene *Rp49* and calculated as 2^-ΔCt^ values, with a detection threshold set at 35 Ct cycles (samples with Ct > 35 were considered uninfected). For each sample. since each sample was prepared by pooling equal RNA amounts from five individuals, we calculated the mean viral load from the five corresponding individual flies measured separately by RT-qPCR (Supplementary File 1).

### Calculating relative aging and pathogenicity

Ratios were calculated by first summarizing the time series of estimated biological age and survival curves (Supplementary File 1) into a single value using the Area Under the Disease Progress Stair (AUDPS) calculation (Simko and Piepho 2012). These values were then used to calculate the relative aging or relative pathogenicity by dividing the AUDPS of the infected samples (either for aging or survival) by the AUDPS of their uninfected controls. The AUDPS calculation was performed using the ‘agricolae’ package (version 1.3-7) in R (version 4.3.2). Survival data was extracted using computer vision assist (automeris.io) from survival plots already published: fly persistent infections come from Castelló-Sanjuán et al. 202510, FHV from Sheffield et al. 2021, DCV-injected male/female from Salminen et al. 2024, DCV-injected with or without *Wolbachia* from Martinez et al. 2019, and OrV from Ashe et al. 2013.

### Statistical analysis

Pearson correlations were calculated in R (v4.3.2) using the *cor.test* function from the ‘stats’ package.

## Funding

Rubén González is supported by an a Pasteur-Roux-Cantarini fellowship of Institut Pasteur. This work was supported by funding from the French Government’s Investissement d’Avenir program, Laboratoire d’Excellence Integrative Biology of Emerging Infectious Diseases (grant ANR-10-LABX-62-IBEID), the Agence Nationale de la Recherche (grant ANR-23-CE15-0038-01, INFINITESIMAL) to Maria-Carla Saleh.

## Author contributions

Conceptualization: R.G. and M.-C.S. Methodology: R.G. Investigation: R.G. and M.C.S, Visualization: R.G. Supervision: M.-C.S. and Writing—original draft: R.G. and M.-C.S. Writing— review and editing: R.G. and M.-C.S. Funding acquisition: M.-C.S. Data curation: R.G. Project administration: R.G. and M.-C.S. Formal analysis: R.G.

## Acknowledgements

We thank members of VIA lab, Artem Babaian, and Allison Bardin for their feedback on the project. Anamarija Butković, Cassandra Koh, and Jared Nigg for their comments on the manuscript. Santiago Elena’s lab for sharing the aligned reads of the *C. elegans* dataset. The authors would like to acknowledge the use of a large language model developed by Anthropic (Claude Sonnet 4) for assistance in refining the R code and editing of this manuscript.

## Data availability

All data genereated in this work is available in Supplementary File 1.

## Supplementary Figures

**Supplementary Figure 1.**
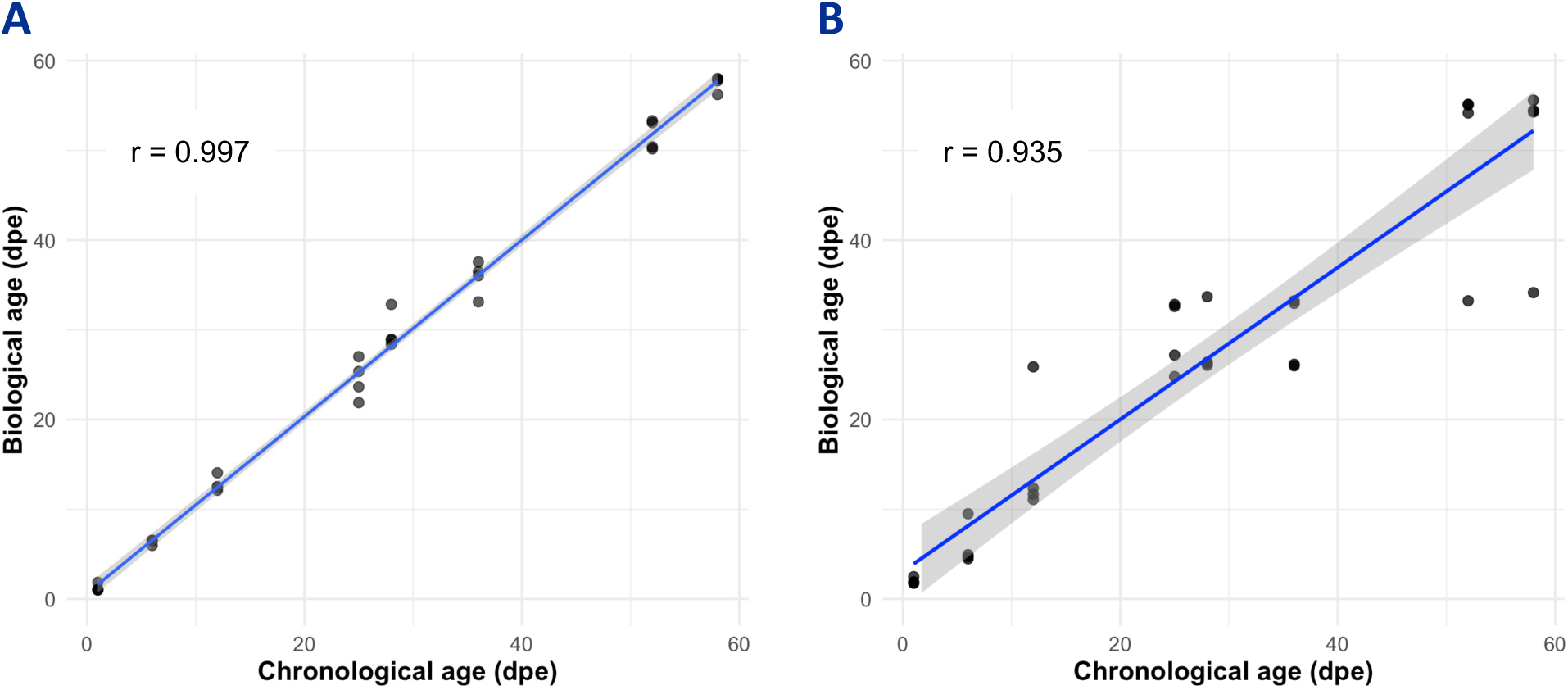
Validation of RAPToR-based biological age estimation in Drosophila reference dataset. **(A)** Full reference performance showing the correlation between chronological age and RAPToR-estimated biological age when all 32 samples (four biological replicates in each one of the eight time points: (1, 6, 12, 25, 28, 36, 52, and 58 days post-eclosion, dpe) are included in both reference construction and age estimation (r = 0.997). **(B)** Leave-one-out cross-validation results showing the correlation between chronological age and RAPToR-estimated biological age when each sample is individually excluded from reference construction and then predicted using the remaining 31 samples (r = 0.935). Each point represents one of the 32 individual predictions, with colors indicating different biological replicates. The blue line represents the linear regression fit with 95% confidence interval (gray shading).

**Supplementary Figure 2.**
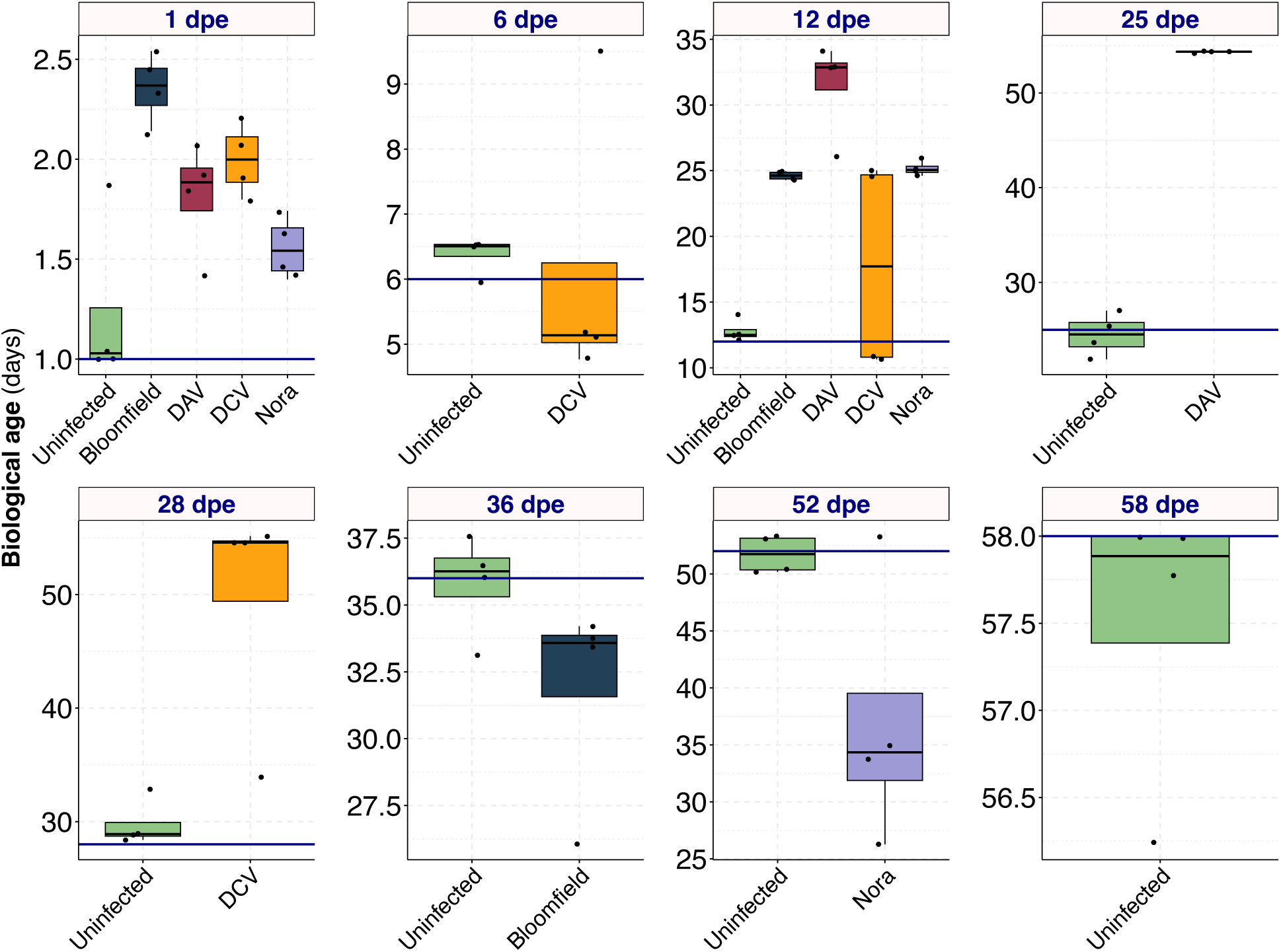
Box plot representation of the biological age of persistently infected and uninfected flies from Figure 2. Each panel represents a specific chronological time point, with the horizontal line indicating the chronological age of the samples.

**Supplementary Figure 3.**
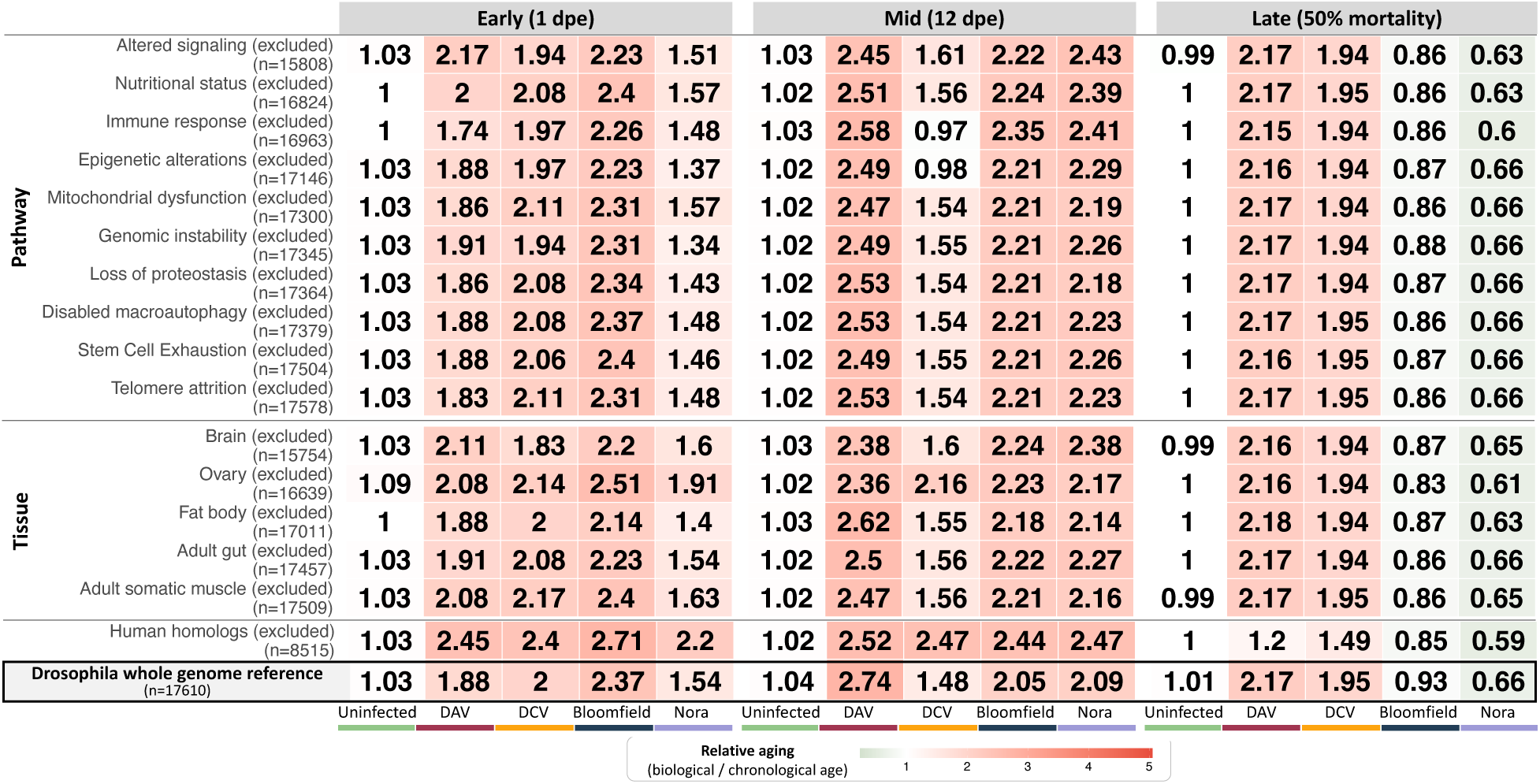
Exclusion analysis. Heatmap shows median relative aging acceleration (biological age/chronological age) using aging references constructed by excluding specific pathway or tissue genes. Early infection (1 dpe), mid-infection (12 dpe), and late infection timepoints (DAV: 25 dpe, DCV: 28 dpe, Bloomfield virus: 36 dpe, Nora virus: 52 dpe, representing ∼50% population mortality). Values >1.0 (red) indicate accelerated aging, values <1.0 (green) indicate slowed aging, and values 1.0 (white) indicate normal aging progression. Numbers in parentheses indicate the number of genes in each reference set. Median absolute deviation values provided in Supplementary File 1.

**Supplementary Figure 4.**
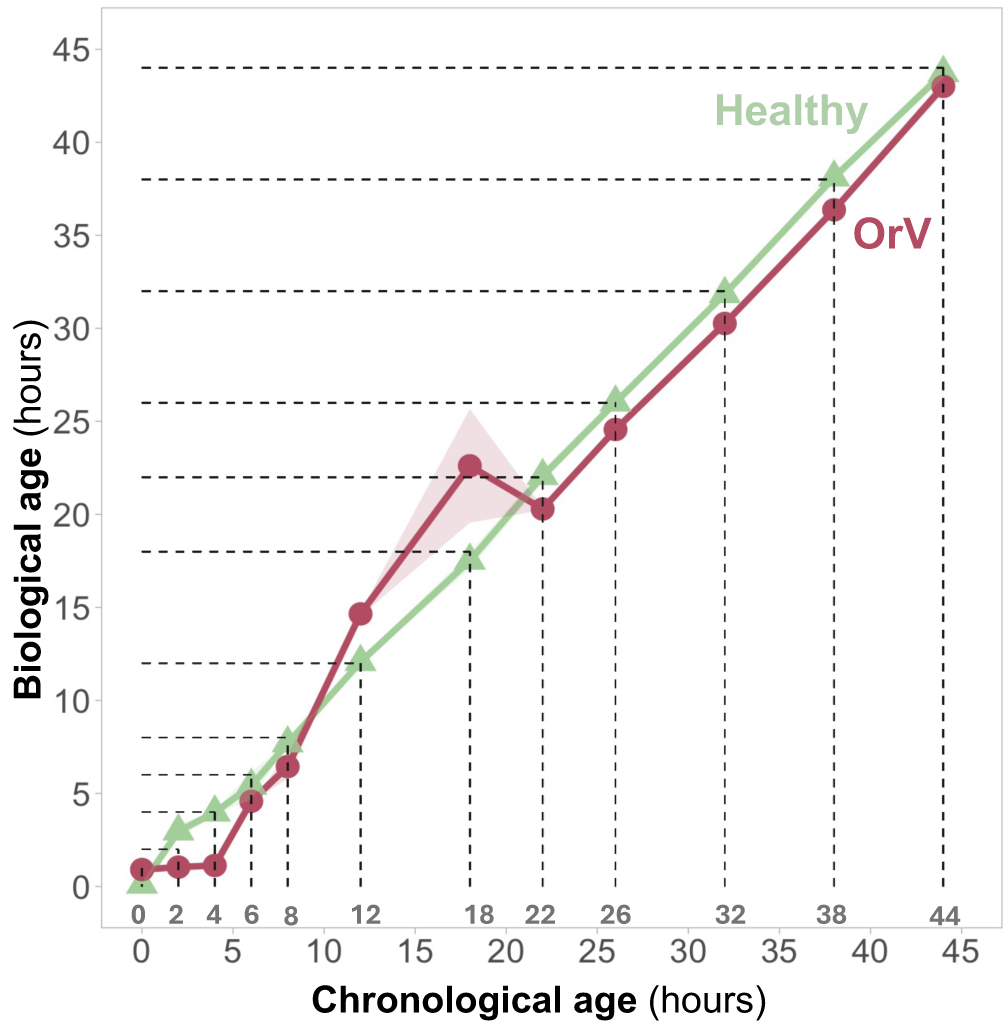
*C. elegans* biological age (Y-axis) at different chronological time points (X-axis) for uninfected animals (green triangles) and those infected with Orsay virus (red circles). Points represent the mean biological age, and the shaded area represents the standard error.

**Supplementary Figure 5.**
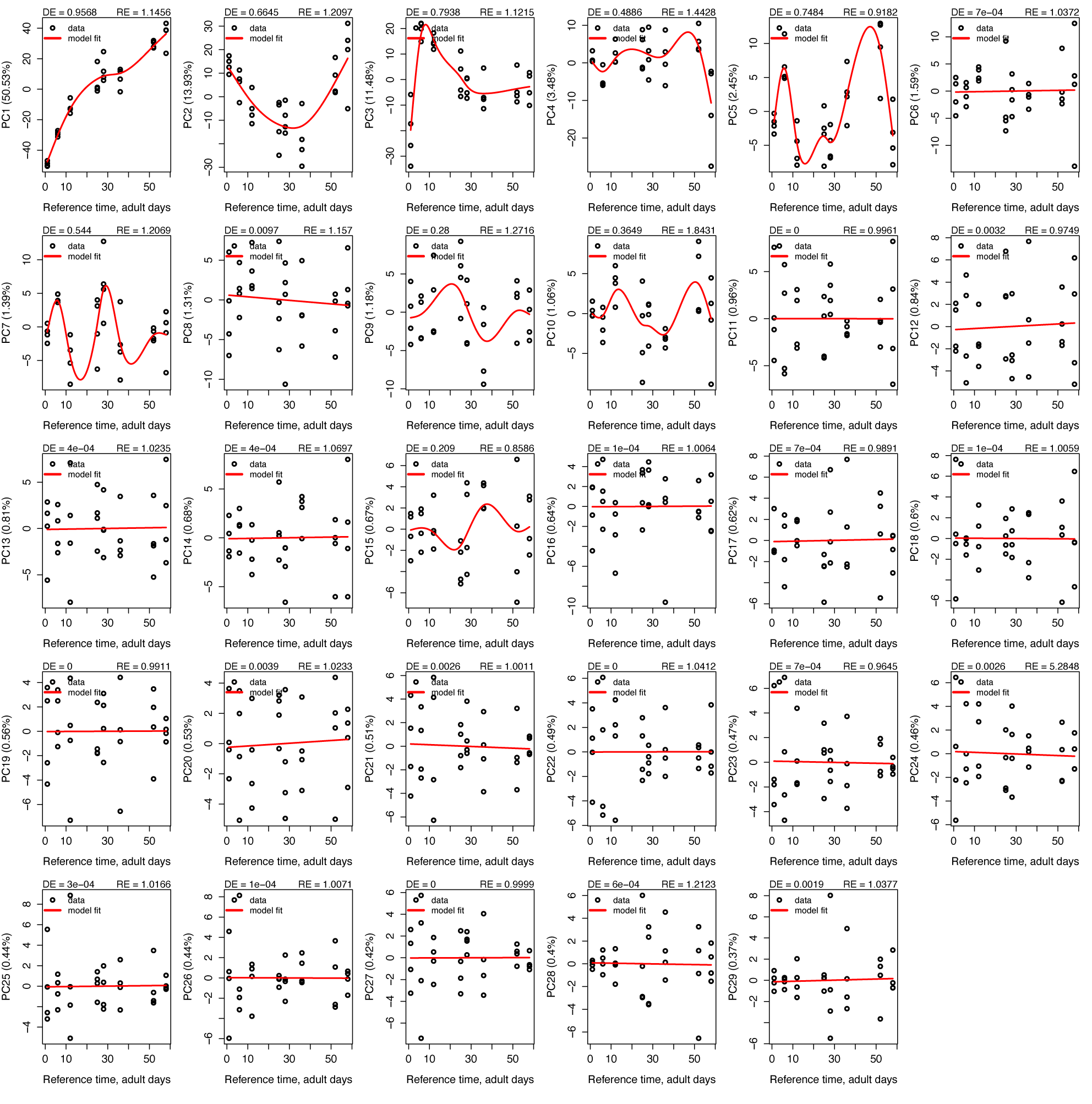
Graphs with the 29 Principal Component Analysis (PCA), which decompose the expression matrix into components, used to create the adult fly reference.

**Supplementary Figure 6.**
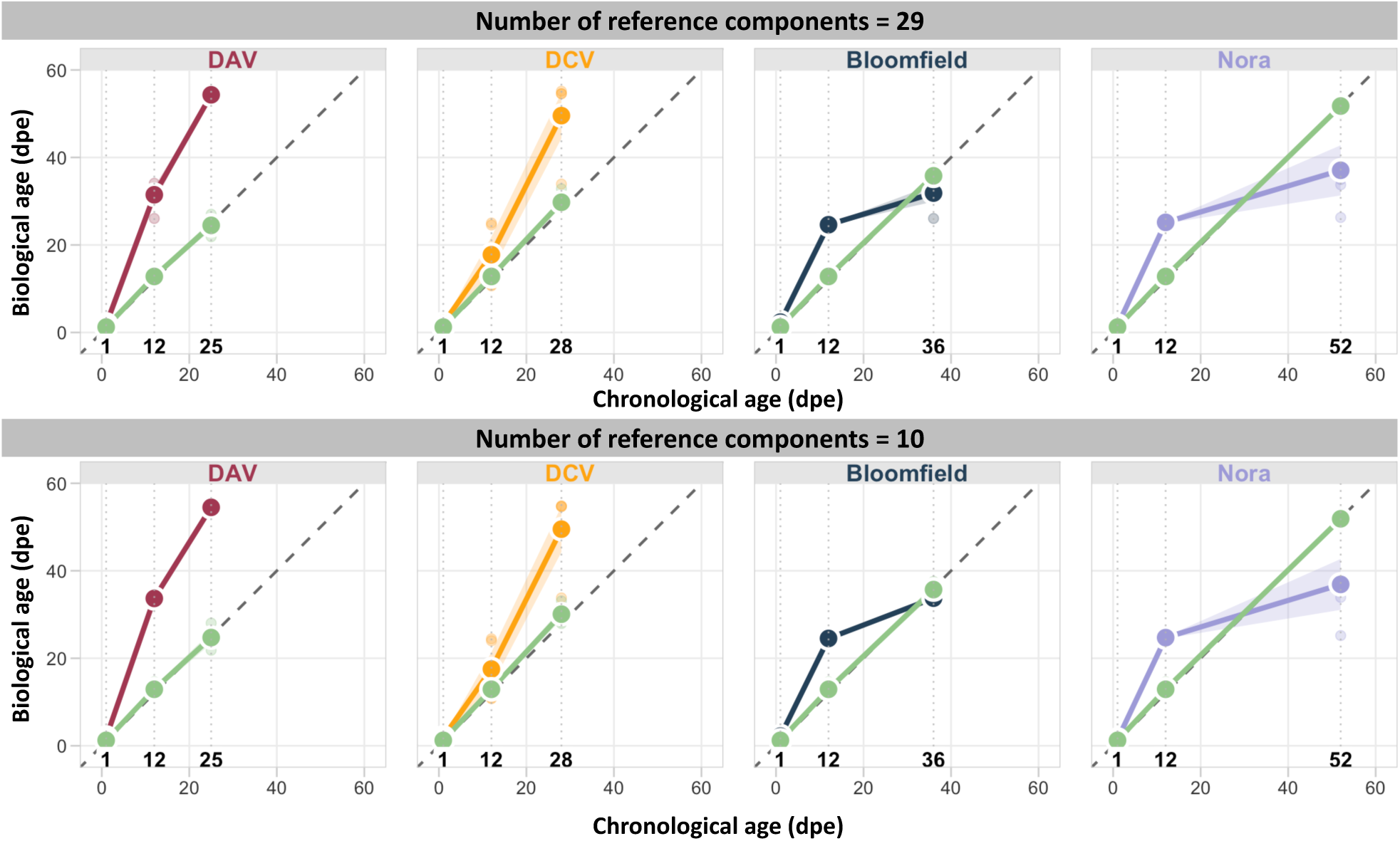
RAPToR biological age estimation is robust to the number of principal components used in reference construction. Comparison of biological age estimates using RAPToR references constructed with either 29 principal components (top panel) or 10 principal components (bottom panel) as a control for overfitting. Each subplot shows the correlation between chronological age and estimated biological age for different viral infections: DAV (red), DCV (orange), Bloomfield virus (blue), and Nora virus (purple), compared to uninfected controls (green). The dashed diagonal line represents perfect prediction (y = x). Both approaches yield highly similar results across all conditions, demonstrating that the aging reference is robust and not subject to overfitting when using the full 29-component model.

## Supplementary Files

**Supplementary File 1.** Data generated per figure.

**Supplementary File 2.** List of pathway and tissue-specific aging genes.

**Supplementary File 3.** List of genes differentially expressed after mating (Newell et al., 2020) excluded from the reference to create a reference for virgin flies.

## Notes

### Competing Interest Statement

The authors have declared no competing interest.

### Summary of Updates

TITLE AND SCOPE: Modified title to emphasize systemic aging effects. Abstract expanded to include pathway analysis, host modulation factors, and persistent aging effects after viral clearance. NEW ANALYSES AND FIGURES: Added pathway-specific and tissue-specific aging analysis examining individual aging hallmarks and organ systems, revealing virus-specific targeting patterns and temporal dynamics and expanding evolutionary conservation analysis using human ortholog genes. (new Figure 3). Included viral load correlation analysis suggesting three distinct mechanisms by which viruses accelerate aging (new Figure 4). Added exclusion analysis confirming systemic rather than pathway-specific effects (new Supplementary Figure 3). ENHANCED CONTENT SECTIONS: Introduction substantially expanded, adding comprehensive literature review on aging biomarkers, computational aging clocks, Drosophila as aging model, and intestinal aging. Results reorganized into six distinct subsections with clearer narrative structure. Discussion expanded. MATERIALS AND METHODS: Enhanced from brief methods to comprehensive section including detailed biological category definitions, pathway and tissue-specific gene set curation protocols, enhanced age estimation methodology, and expanded statistical analysis descriptions. Added complete gene lists for aging hallmarks curated from established frameworks and FlyBase annotations. Added cross-validation analysis for aging clock validation.

